# Estimating likelihood from simulations in bayesian inference of ecological models with historical contingency

**DOI:** 10.1101/453027

**Authors:** Stéphane Dupas

## Abstract

Ecological patterns result from historical contingency and deterministic processes. Taking apart these processes to extract probabilistic models of ecological dynamics is of major importance for ecological forecasting. Due to the high dimensionality of historical contingency it is usually difficult to sample history from observed patterns. In environmental population genetics, the number of possible genealogies linking genetic data and environemental data through demographic and niche models is almost infinite. In ecosystem dynamics time series, the patterns are determined as much by probabilitic model parameters, as by historical variables contingency. Aproximate bayesian computation allows to use simulations to aproximate this inference process. The rationale is to simulate data, extract summary statistics and retain in the posterior, the parameters values that produced simulations with summary statistic close to observed summary statistics. The major drawbacks of this approach is that summary statistics distance is not exhaustive regarding model likelihood and may biais the results. In the present work, I show that if we can simulate the historical contingency from observed data and probabilistic model in a backward approach, we can to use the simulations to estimate a pseudo likelihood that can be used in bayesian inference. I apply to genealogy sampling in environmental demogenetics and ecosystem dynamics modelling.

## Introduction

Bayesian inference is a powerful approach to combine prior information to field data and improve models instead of rejecting them. However complex model bayesian sampling can become rapidly intractable, particularly for biological and social processes in which historical contingency plays a major role. Aproximate bayesian computation has become a popular solution for sampling historical processes in ecology [1]. The parameters filtering is obtained through the selection of simulations that are the closest to the observations. However selecting simulations brings back to the problem of defining summary statistics that represent the probabilistic model. Simulation and particle filtering using MCMC have been proposed for dynamic bayesian network [2]. Here I propose to use simulation to complete data in order to estime an aproximate likelihood forbayesian inference. I claim that the best summary statistics would be the probability to observe altogether data and simulations knowing model parameters. In many case this probability can be calculated. The only condition is to be able to formulate a probabilistic model to simulate history from observed data and model. For instance, it is possible to calculate the probability of a genealogy, knowing (1) genetic distributions in allelic and geographic space, (2) environemental data and (3) an environemental demogenetic model including mutation model, niche model and dispersion model. If we average this probability over genealogical simulations we get an estimate proportional to the probability of the parameter set.

## Model

### Probability and simulation of hidden history knowing indicator data and ecosytem model

We divide the model *M* into the sampling model *M_S_* that link ecosystem variable history *H* to data *D*, and the ecosystem history model *M_H_* that drive ecosystem *H* variables dynamics (*M* = {*M_H_, M_S_*}). Each has its set of parameters *θ_S_* and *θ_H_*. We know how to calculate *p*(*D/H*, *M_S_*) and *p*(*H*/*M_H_*). The probability of a given history knowing data and model is given from bayesian rule:

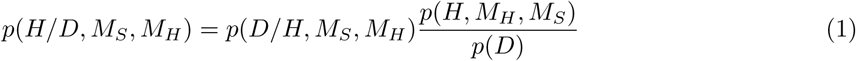

Sampling model *M_S_* and ecosystem model *M_H_* are independent processes : sampling of data from *M_S_* and *H*, is independent from sampling of history *H* from ecosystem model *M_H_*. We then have: *p*(*D*/*H*, *M_S_*, *M_H_*) = *p*(*D*/*H*, *M_H_*) and *p*(*H*, *M_H_*, *M_S_*) = *p*(*H*, *M_H_*) × *p*(*M_S_*). Since we have also *p*(*H*, *M_H_*) = *p*(*H*/*M_H_*) × *p*(*M_H_*), equation (1) gives:

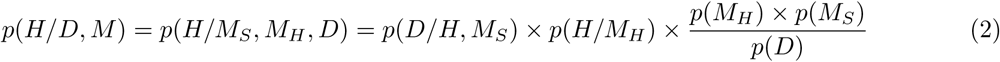

From this distribution, we can simulate a history *H_i_* by using prior information on *M_H_*, *M_S_*, and *D*. *D* is independent on model parameters. We also have the probability of each historical simulation *H_i_* knowing data and model using (2).

### Estimation of model likelihood by averaging probability of histories

Now we can estimate the probability of the data knowing the model. We consider [*H*] = *H_i_*, *i* ∈ [1‥*n*] the set of all possible histories.

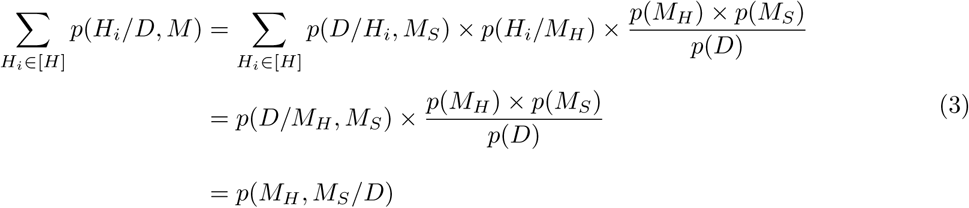

We see in the equation that the sum of data probability knowing sampling model from history multiplied by history probabilities knowing ecosystem model plays the role of a likelihood function in the classic bayesian equation.

We can have an estimation of the pseudo likelihood 
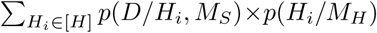
 by simulating a number of times *Hi* using distribution in (2) and averaging the pseudo likelihood. Since the simulation will produce the most probable histories, after a number of simulations, we will get an estimate proportional to the probability of all histories.

We can therefore name this method bayesian inference with aproximated likelihood (BIWAL).

I now apply to ecosystem and coalescent models.

#### Algorithm

The following algorithm will be used :

1. Set sampling and ecosystem models *p*(*D/H, M_S_*) and *p*(*H/M_H_*)
2. Set sampling and ecosystem model parameters priors *p*(*θ*), where *θ* = [*θ_S_, θ_H_*], and *θ_S_*, and *θ_H_* are the *M_S_* and *M_H_* model parameters, respectively.
3. Set Metropolis sampling generation *g* = 1, a parameter proposal rule from *θ* to *θ′*, and a *thining* value for sampling frequency of *θ*
4. Sample *θ_g_* from prior and calculate posterior *θ_g_* the same way as described in steps 6 and 7 for *θ′*
5. Propose a *θ′* from *θ* using metropolis proposal rule
6. Repeat for i *∈* 1 : *n*

a. Simulate History *H_i_− > p*(*H_i_/M, D*)
b. Calculate *p*(*H_i_/M, D*)
7. calculate posterior *p*(*θ′/D*) from equation (8)
8. if *p*(*θ′*) > *p*(*θ*) set *θ* = *θ′* if *p*(*θ′*) <= *p*(*θ*) set *θ* = *θ′* with probability 
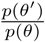
9. save *θ_g_* if *g* is a multiple of *thining*
10. *g* = *g* + 1 go to step 5.

### Application to coalescent model

In Dupas (unpublished), we developped an algorithm to simulate and estimate the probability of a genealogy of individuals knowing their spatial and genetic distributions, *D* and *G*, respectively, the environmental data *E*, and the environmental demogenetic model *M_D_, M_G_* (equation 13, Dupas 2018). Consider a simulated genealogy *H_i_* = *genealogy_i_* sampled from *p*(*genalogy/M_D_, M_G_, G, D, E*) then according to (6)

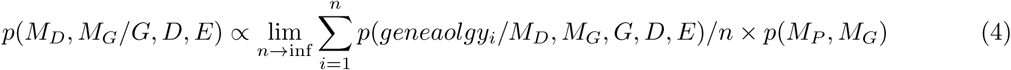

### Application to crop yield prediction

We develop here a crop yield model we apply the growth rate function based on existing apriori deterministic model constructed from literature information [3].

1. *D_nr,i,j_* = number of rainy days in month *i*, year *j*
2. *D_P r,i,j_* = precipitation
3. *D_T,i,j_* = average daily temperature
4. *D_T x,i,j_* = maximum daily temperature
5. *D_T n,i,j_* = minimum daily temperature
6. *D_ET P,i,j_*] = potential evapotranspiration
7. *D*_*Y*,10,*j*_ = observed maize yield at harvest
8. *H*_*Y*,10,*j*_ = expected maize yield at harvest
9. *H_M,i,j_* = plant mass
10. *H_vgr,j_* = Maize variety maximum growth rate
11. *H_vprd,j_* = Maize variety potential resistance to drought (decrease in yield with respect to per unit decrease in ETa) [3]
12. *H_ET A,i,j_* = realized evapotranspiration

In addition we consider *θ_T gr_* the temperature coeffient of growth rate. We dispose of monthly data for each of these historical variables, for the 94 French departments for 38 years. In this example there is no missing data.

The ecosystem model is as follows :

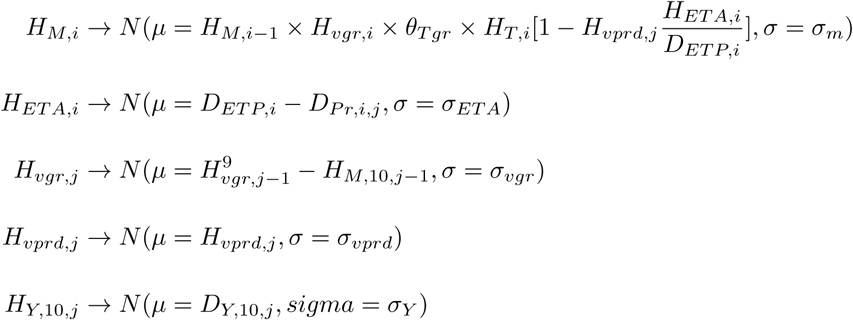

## Discussion

I propose a new method to sample complex ecological models with important historical contingency. The rationale is to average historical processes calculating simulations probability and to sample parameters using Metropolis Hasting algorithm. The requisite is to be able inverse part of the model using bayesian rules in order to simulate history from data. This backward approach is classic in populations genetics where coalescent theory has been developped since the 1980’s [4]. But similar models can be developped in other area in ecology. In particular for ecosystem dynamics inference from time series data [5].

## Acknowledgments

This work has been supported by IRD EGCE research unit project and BASC labex

